# Exploring the functionality of market-available tools for neural recording

**DOI:** 10.64898/2026.06.20.720337

**Authors:** Khashayar Esmaeilzadeh, Mohammad Hosseini, Seyed Ahmad Etghani, Abdolhossein Vahabie, Milad Yekani

## Abstract

Low-cost and open-source neural recording systems are increasingly important for expanding access to electrophysiological research. However, many existing platforms still rely on specialized hardware or limited modularity, restricting flexibility for laboratories seeking customizable solutions. Here, we developed and evaluated a modular neural recording platform constructed entirely from commercially available components. Recordings were compared against the ground truth. The platform successfully recovered local field potential (LFP)-like waveforms in most conditions and detected spike-like activity during direct connection recordings. Principal component analysis and k-means clustering further demonstrated the ability to distinguish multiple simulated spike waveforms. Signal quality varied across configurations, with saline recordings and preamplifier integration introducing increased noise and reduced detectability. These findings demonstrate the feasibility of building affordable and modular electrophysiology systems using widely accessible hardware. Although the current implementation has limitations in sampling rate, noise performance, and in vivo validation, the presented framework provides a practical foundation for future customizable open-source neural recording.

## Introduction

Electrophysiology has been one of the major techniques for studying the brain and the primary method for providing signals in brain-computer interfaces. With the growth of the neuroscience field in recent decades and the growing desire to connect devices to our brains, there is a considerable demand for the development of brain recording devices. Conventional laboratory recording devices are usually expensive and limited in customization. These barriers motivate researchers to seek open-source, cost-effective software and hardware solutions. Some attempts have been made in this direction, but with the advent of technology, more affordable solutions are becoming available, and there is always room to make them even cheaper. One approach to achieving this goal is to identify existing market solutions and test their applicability.

Open-source neurotechnology initiatives demonstrate that accessible hardware can successfully support meaningful scientific work. For example, OpenBCI has played a notable role in popularizing low-cost biosignal acquisition by providing community-supported EEG/EMG solutions (Laport et al., 2019; Niforatos et al., 2025). However, such platforms are not built primarily from readily available off-the-shelf components. As a result, researchers seeking to design adaptable system models often encounter challenges (Erofeev et al., 2023).

For laboratories that rely on modularity and hardware flexibility—for example, it becomes essential to systematically evaluate market-available components such as analog front-ends, microcontrollers, and discrete ADCs. Recent open-hardware platforms have shown that repurposing commercially available electronic parts can be an effective strategy for lowering cost while maintaining adequate data quality for neural experimentation (Voitiuk et al., 2021). A modular device—where preamplifiers, cables, digitizers, and microcontrollers can be replaced or upgraded independently—creates opportunities for rapid iteration and community-driven improvement, ultimately increasing reproducibility across research groups.

In response to these needs, we developed and evaluated a low-cost, modular neural-recording system constructed entirely from widely available hardware. The platform combines a lightweight, small-animal-compatible headstage preamplifier with a digitalization pipeline using the high-resolution ADS1251 ADC interfaced with an STM32 microcontroller. The design emphasizes a balance between noise immunity, low weight, low power consumption, and affordability—key requirements for electrophysiological experiments in small animal models. All circuit schematics, PCB layouts, and control software are openly provided to encourage modification and adoption by the neuroscience community.

By benchmarking this system through simulated and biological recordings—including direct electronic ground-truth testing, saline-bath evaluation—we assess the feasibility and limitations of using low-cost commercial components for neural data acquisition. Our results demonstrate both the potential and current constraints of this modular approach and outline a path forward for developing future customizable and fully open neural-recording devices.

## Methods

### Preamplifier design

A headstage preamplifier using AD623 was designed and assembled by our team. The default schematic and PCB for the preamplifier include a low-pass filter, which can be replaced with a high-pass filter with respect to the intended recording (LFP or spike). PCBs and schematics were designed via Altium Designer version 25.0.2 (Built 28).

### Method of digitalization

Two different digitalization methods were compared in this work. First was through the Open-BCI Cyton board V3.3, which uses ADS1299 as its analog front-end. Second, we utilized ADS1251 readings through an STM32 board, which we programmed.

### Computer interface

#### ADS1251

To fully utilize the high sampling rate of the ADS1251, we implemented data acquisition on an STM32 microcontroller instead of Arduino boards. The STM32 reads the ADC output at high speed and streams the data through a serial connection at 1 Mbps. On the computer side, a custom Python GUI (based on Tkinter) was developed to handle the incoming serial data. The interface supports real-time monitoring by plotting the most recent 200 samples dynamically, while also storing the entire dataset for later analysis. Additionally, users can save the recorded signals in CSV format, manually send commands to the STM32, and control acquisition (start/stop) through the graphical interface. As the STM32 streams ASCII, newline-terminated sample values over a virtual COM port at 1,000,000 baud, raw data can be inspected directly using a serial terminal (e.g., PuTTY) without the Python GUI.

### Hardware testing

We used a BlackRock Digital Neural Signal Simulator as our primary testing measurand, both by direct connection and through a saline solution (fig. 1).

**Fig. 1.**
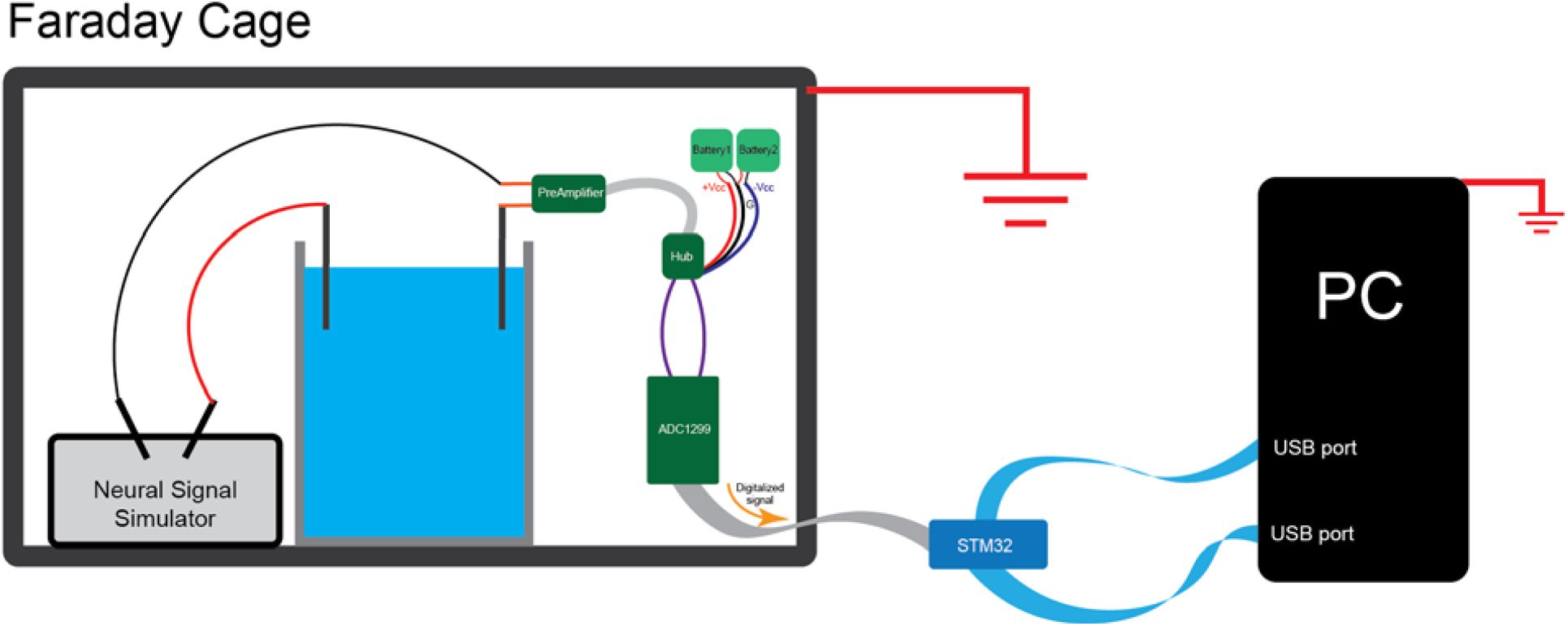
Illustration of the hardware testing setup. The recording device contains modular stages (preamplifier, hub, ADC, STM32), which were tested both with and without the preamplifier. For raw signals, we used a BlackRock Digital Neural Signal Simulator. Both a saline connection, as presented in this figure, and a direct connection to the simulator were recorded. The setup allows for a USB input to a computer.

## Results

Direct Connection and Saline tests of the ADC (with or without the pre-amplifier) were recorded in a Faraday cage, and subsequently compared and contrasted in Fig. 2. For a ground truth (GT) signal, we used data from a BlackRock Amplifier Manifold (gain set to 1) and a BlackRock NeuroPort Neural Signal Processor, connected directly to the Simulator (without saline) (fs=30kHz). BlackRock processed signals were converted to MATLAB-readable format through the NPMK toolbox (Blackrock Neurotech, 2015/2025). As the saline bath and preamplifier set a different amplitude scale, GT recordings were proportionally y-axis scaled for visual comparison in Fig. 2b-d.

**Fig. 2.**
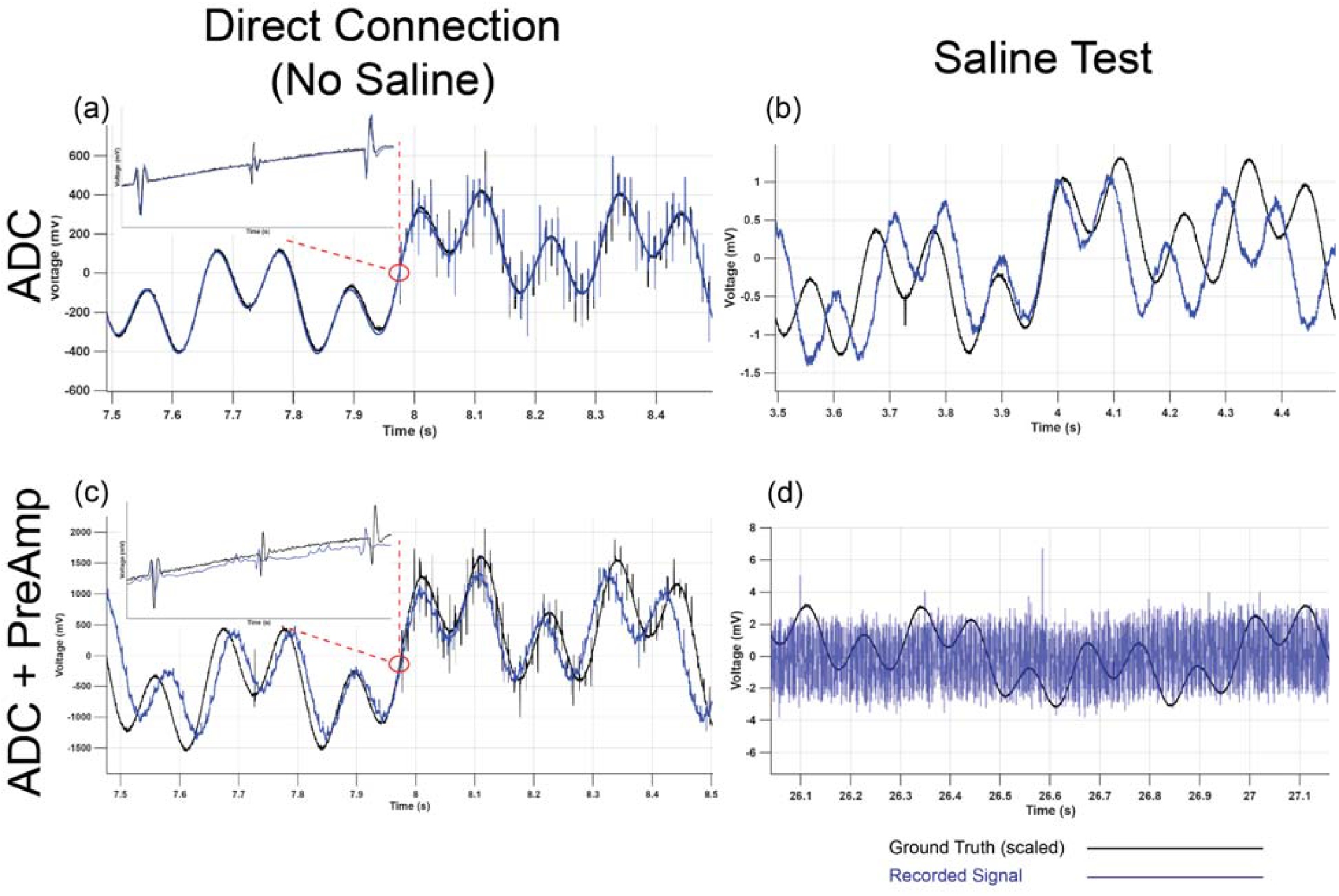
Comparative plots of the recorded signals. The ground truth was established using recordings from a BlackRock Amplifier Manifold (gain set to 1) and a BlackRock NeuroPort Neural Signal Processor, connected directly to the simulator (without saline) (fs=30kHz).

The sampling frequency of our recording device was 11110.601 Hz, recorded at a baud rate of 1000000 through the USB port. Traces were converted from 24-bit ADC codes to microvolts-seconds, median-detrended (4.0 s window), and zero-phase FIR high-pass filtered at 0.5 Hz (FIR order capped at 20,000). A strong 50 Hz notch (Q=4) was optionally applied twice for comparison. For the Saline, ADC (Fig. 2b), a moving average of 5ms was applied.

Our results in direct connection visually reached the waveform (Fig. 2a) of the LFP signal of our GT, with residual (GT vs Device), and peak/RMS signal-to-noise (SNR) values of 10.70 dB and 8.93 dB, respectively, for the LFP band of 1 to 300 Hz. With the introduction of the preamplifier module (Fig. 2c), the SNR values changed to 5.28 dB and 9.47 dB, respectively. Moreover, spikes were visually and quantifiably detectable (ADC: threshold 5mV=1.26σ, ADC+PreAmp: threshold 200mV=5.86σ) in our direct connection recordings, with an average of 8 data points per spike.

For the saline test without the preamplifier (Fig.2. b), an LFP was detected with residual and peak/RMS SNRs of 2.41 dB and 9.67 dB. Adding the preamplifier in saline resulted in no detectable signal in our recording (Fig. 2c). For both saline tests, no spikes were able to be detected.

As a further analysis, as the BlackRock Simulator generated three distinct waveforms for spikes, a PCA (n=448) was calculated on the snippets of 5 samples pre- and post-threshold crossing (Fig. 3a). The first 6 principal components, explaining 97.2% of the total waveform variance, were picked for k-means (k=3) clustering (Fig. 3b). The analysis appears to have successfully recovered the three known units, with cluster quality quantified via peak signal-to-noise ratios of [11.0790 7.7347 10.1051] (linear) equivalent to [20.8900 17.7688 20.0908] dB, contrasted to 54229 samples from baseline.

**Fig. 3.**
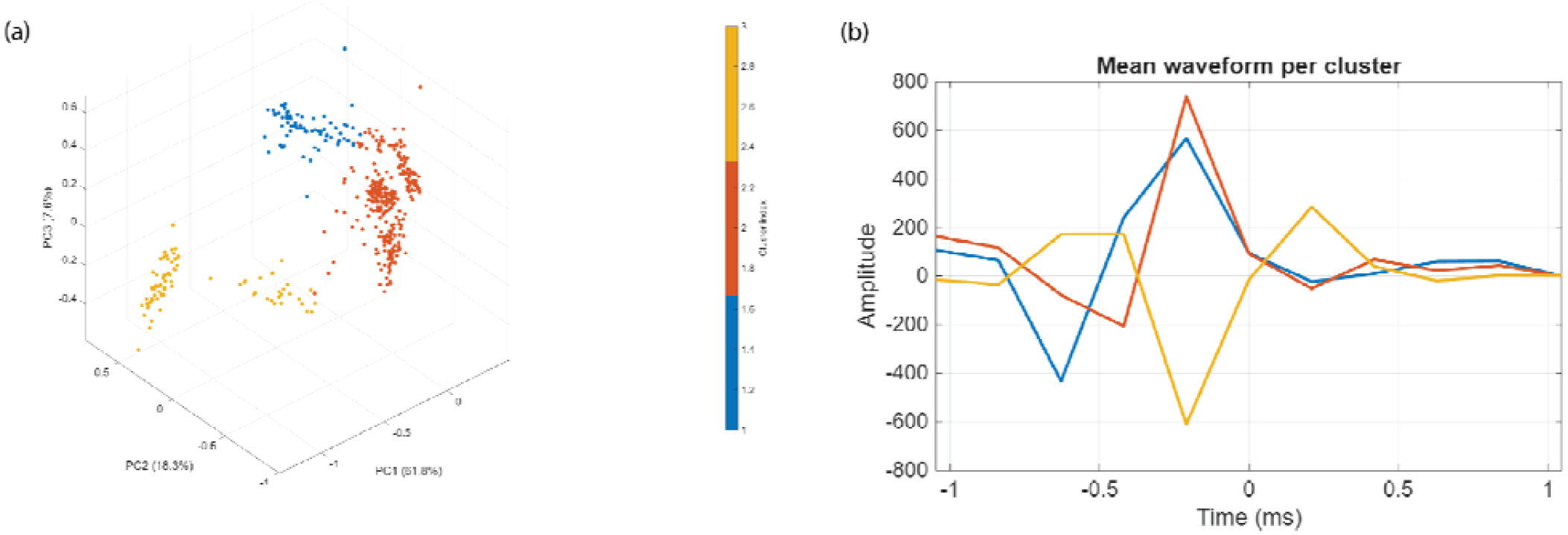
PCA-based spike sorting on direct connection, ADC+preamplifier recording. (a) Principal component analysis of 448 detected spikes (10-sample snippets: 5 pre- and post-threshold) visualized in 3PC space (PC1=61.8%, PC2=18.3%, PC3=7.6%). (b) Mean peak-aligned waveforms by k-means (k=3) automatic clustering.

## Discussions

We carried over an approach to build and test a neural recording device through a low-cost protocol. The ADS1256 was selected as the digitalization method, and an STM32 operating code was developed to interface the ADS with a computer via the USB port. A real-time plotter for the data was built using Python. In addition, a low-weight headstage preamplifier was printed, assembled, and tested with the ADS (Fig. 1). For evaluation, recorded LFP and spike signals were compared with standard BlackRock-processed ones as the ground truth. We compared the performance of the ADS with and without the preamplifier, by direct connection and saline bath (Fig. 2. a-d). Spikes were recoverable in the direct connection results (Fig. 2a & 2c), and the LFP waveform was recoverable in all tests except ADS+Preamp in saline bath (Fig. 2d). In a further step, we tried to recover three distinct spike waveforms generated by the simulator in the direct connection of the ADS+Preamplifier (Fig 3). With a k-means (k=3) autoclustering, the waveforms appear to be successfully distinguished in a peak-aligned plot (Fig. 3b).

There are notable open-source initiatives helping scientists afford to use electrophysiology hardware. Some examples are OpenBCI (Laport et al., 2019), NeuroRighter (Rolston et al., 2009), and OpenEphys (Siegle et al., 2017). But they often rely on specialized components, complex fabrication, and firmware/software stacks. Thus, there is room for improvement in the availability and affordability of electrophysiology recording devices. We demonstrate a feasible pathway to building a low-cost neural recording device using off-the-shelf components (the ADS1256 and an STM32 microcontroller). Importantly, the system is designed as a modular platform that serves as a practical foundation for more advanced neural recording devices. While not intended as a finalized solution, its flexible architecture allows future development toward higher performance, additional functionalities, and more sophisticated experimental applications.

The reached sampling rate raises problems for spike recordings. For spike sorting and detection, typically the sampling rate is around 25kHz or more (Navajas et al., 2014). High frequency components of spikes could be up to Due to the Nyquist theorem, high frequency components of spikes (>5555.3Hz) may not be detected via this sampling rate. Although the waveforms appear to be successfully distinguished (Fig. 2b), the resolution might not be enough for spike sortings in a hands-on lab setting for animal models. The saline tests suggest that the preamplifier module could render ineffective for in-vivo electrophysiology, as no LFP signal appeared to be detectable in the recordings. Moreover, for an LFP recording session, although the waveforms are distinguishable in the saline test, it is worth noting that it comes with a high noise (2.41 dB residual SNR) as a caveat. Another downside of the current chosen modules is the skewed signals, which appear to be stretched compared to the ground truth in Fig. 2b,c. The detectable noise may render the current assemblages of the modules insufficient for an awake, behaving animal LFP recording. Furthermore, our evaluation could fall short as it lacked in vivo recording sessions.

More robust connections at and between different modules are possible for protection against noise and artifacts, for instance, by using coaxial cables or employing Electromagnetic Compatibility (EMC) testing to mitigate electromagnetic interference. For the headstage, we plan to design a micro version of the ADS board to print and test for LFP recording. Other computer interface modules, like Arduino boards, could be tested for more accessibility of the method for neuroscience researchers. Other possible developments include working on a wireless module for communication between a headstage and the computer interface instead of a tethered hub. For the evaluation stage, recording from anesthetized or behaving animals through established protocols could be performed, following saline tests. An example of using anesthetic signals is presented at (Abolghasemi et al., 2024).

## Conclusion

Based on our findings, this simple, highly cost-effective hardware setup provides a practical framework for developing brain signal recording devices. Future work can build on this foundation to develop more complex, application-specific systems tailored to diverse experimental requirements.

## Notes

### Competing Interest Statement

The authors have declared no competing interest.

